# Non-fibrillar prion protein oligomers transmit structural information during early assembly

**DOI:** 10.64898/2026.03.23.713466

**Authors:** Stephanie Prigent, Ariane Deniset-Besseau, Jeremi Mathurin, Angélique Igel, Hannah Klute, Jan Bohl, Guillaume van der Rest, Christian Malosse, Sophie Lecomte, Hadi Rezaei, Joan Torrent, Vincent Béringue, Alexandre Dazzi, Davy Martin, Human Rezaei

## Abstract

The prion paradigm is founded on the transmission of structural information from aggregated assemblies to soluble protein substrates, a process classically attributed to fibril-end-mediated templating. Whether non-fibrillar assemblies or transient oligomeric states can also participate in folding information transfer remains unclear. Here, using recombinant prion protein (PrP), rational mutagenesis, hetero-oligomerization assays, arrested reaction conditions, and single-particle atomic force microscopy coupled to infrared nano-spectroscopy (AFM-IR), we examine the earliest stages of PrP assembly from the perspective of folding information transmission. We show that polymerization-defective PrP variants can be incorporated into oligomeric assemblies through structural complementation, in which folding information supplied by wild-type PrP restores their assembly competence. Under arrested reaction conditions, transient polymerization-competent conformers further contribute to folding information transfer at subcritical concentrations. Domain-resolved analyses reveal a modular oligomeric architecture in which a β-sheet-rich B domain constitutes the primary scaffold for folding information transfer. Preformed O1 oligomers then act as autonomous conformational templates that promote mutant incorporation and undergo hierarchical condensation through accretion of a structurally distinct E domain. Together, these findings demonstrate that non-fibrillar PrP oligomers and transient assembly intermediates can store and transmit folding information and may function as oligomer-based secondary nucleation platforms, expanding the conceptual framework of prion assembly beyond fibril-end elongation alone.

## INTRODUCTION

In mammals, prion diseases refer to a class of infectious neurodegenerative pathologies in which the infectious conformer called PrP^Sc^ acts as template transferring a misfolding information into the physiological conformer denoted PrP^C^. The process is commonly called templating process. This simplified vision where the couple PrP^Sc^ and PrP^C^ are assimilated to respectively donor and receiver of a structural information constitutes the bedrock of prion theory (1).

From a folding perspective, the templated conversion of PrPC into PrPSc is fundamentally distinct from most protein misfolding disorders in which the prion paradigm has been applied. In fungal prions (Ure2, Sup35, Het-s) and prion-like proteins, as well as in amyloid-β and α-synuclein, the aggregation-prone regions are intrinsically disordered (2–4). In these systems, templating primarily induces folding of a disordered substrate into an ordered amyloid state, such that the transmitted information is essentially a folding instruction (5). Mammalian prions differ in that PrPC is a compact, predominantly α-helical protein with high thermodynamic stability (6, 7). Its conversion therefore cannot be reduced to a simple folding reaction. Instead, templating must involve at least two coupled steps: (i) partial unfolding that destabilizes the native α-helical architecture (8), followed by (ii) refolding into a β-rich PrPSc like conformation imposed by the templating assembly (i.e., PrPSc aggregates) (9). How these unfolding and refolding events are orchestrated, and how structural information is transmitted during this process, remain largely unresolved.

Despite this complexity, prion templating is generally assumed to be restricted to fibrillar assemblies. In this canonical view, fibril extremities act as the active sites of conversion, where unsatisfied hydrogen bonds and exposed hydrophobic clusters (10, 11) catalyse conformational change through an induced-fit mechanism (12).While this model readily explains amyloid elongation, it provides little insight into the contribution of oligomeric species during the earliest stages of assembly. The role of oligomers in prion aggregation remains poorly defined, and their ability to initiate polymerization or transmit structural information is largely unexplored. Crucially, protofibrils, often assimilated to oligomers, and fibril fragments must be distinguished from *bona fide* oligomers (13–15). The former are on-pathway species that are structurally quasi-equivalent to mature fibrils and efficiently promote aggregation through classical templating or seeding mechanisms. In contrast, off-pathway oligomers are structurally distinct from fibrils, and whether they can store or propagate conformational information independently of fibril elongation remains an open question(16–18).

One such assembly, termed O1, is a discrete, non-fibrillar PrP oligomer that can be reproducibly generated and isolated, providing a well-defined system to examine whether off-pathway assemblies can store and transmit structural information independently of canonical fibril elongation (6–8). O1 is formed upon thermal unfolding of full-length recombinant sheep PrP and corresponds to a soluble oligomeric population enriched in β-sheet structure following partial destabilization of the native α-helical fold (7, 8). Structural and kinetic analyses indicate that O1 arises from a parallel oligomerization pathway that is largely disconnected from the nucleation-elongation mechanism leading to amyloid fibrils, although secondary worm-like supramolecular assemblies can be derived from the condensation of O1 under specific conditions (8). Formation of O1 involves an early expansion of the C-terminal globular domain and critically depends on the H2-H3 helical region, which constitutes the minimal oligomerization domain of PrP (19–21). Despite its non-fibrillar nature, O1 displays pronounced biological activity, as recombinant PrP oligomers of this class are neurotoxic both in vitro and in vivo, in sharp contrast to monomeric PrP and mature fibrillar assemblies (22, 23). Together, these properties identify O1 as a structurally and biologically relevant oligomeric state and a tractable model to investigate the mechanisms of prion conversion beyond fibril-end-mediated templating.

In this work, we investigate the earliest stages of prion protein assembly to determine whether structural information transfer can occur outside the classical framework of fibril-end elongation. By combining rationally designed PrP mutants, hetero-oligomerization assays, and arrested reaction conditions with single-particle high-resolution liquid atomic force microscopy and infrared nanospectroscopy (AFM-IR), we dissect how folding information is transmitted during the initiation of oligomer formation. This integrated approach allows us to distinguish complementation mediated by polymerization-competent PrP variants, contributions from transient folding intermediates, and templating by discrete non-fibrillar oligomeric assemblies. Through domain-resolved structural analysis, we further examine how oligomeric scaffolds encode and propagate folding information, providing a framework to assess how non-fibrillar quaternary structures can initiate and organize prion assembly independently of fibril elongation.

## RESULTS

### Structural Complementation and Folding Information Transfer During PrP Oligomerization

To investigate whether structural information transfer can occur during non-fibrillar PrP oligomerization, we implemented a structural complementation strategy based on hetero-oligomerization. Single–amino acid substitutions were introduced to selectively impair PrP polymerization, and Hetero-oligomerization assays were used to determine whether interaction with polymerization-competent wt PrP during oligomerization could restore the missing folding information required for assembly.

To implement this strategy, we generated a panel of full-length PrP variants carrying single–amino acid substitutions within helices H2 (residues 188–191) and H3 (residues 206–209), regions previously shown to govern PrP oligomerization and pathway selection. Alanine substitutions within H2, including K188A and H190A, selectively promoted oligomerization through the O1 pathway, yielding robust O1 populations under conditions permissive for wild-type PrP assembly. In contrast, substitutions within H3, notably I206A and I208A, markedly impaired PrP self-oligomerization, resulting in variants that remained predominantly monomeric and failed to generate detectable oligomeric species under the same conditions (Fig.1A and B). These polymerization-defective variants provided a stringent readout for complementation and structural information transfer. If oligomerization involves structural complementation, incorporation of such mutants into oligomeric assemblies should restore the missing folding information required for productive oligomer growth. Conversely, failure of the mutant to adopt a quasi-equivalent fold upon interaction with wild-type PrP would result either in its exclusion from the oligomeric assembly or in dominant-negative (“suicide”) inhibition of oligomerization.

**Figure 1:**
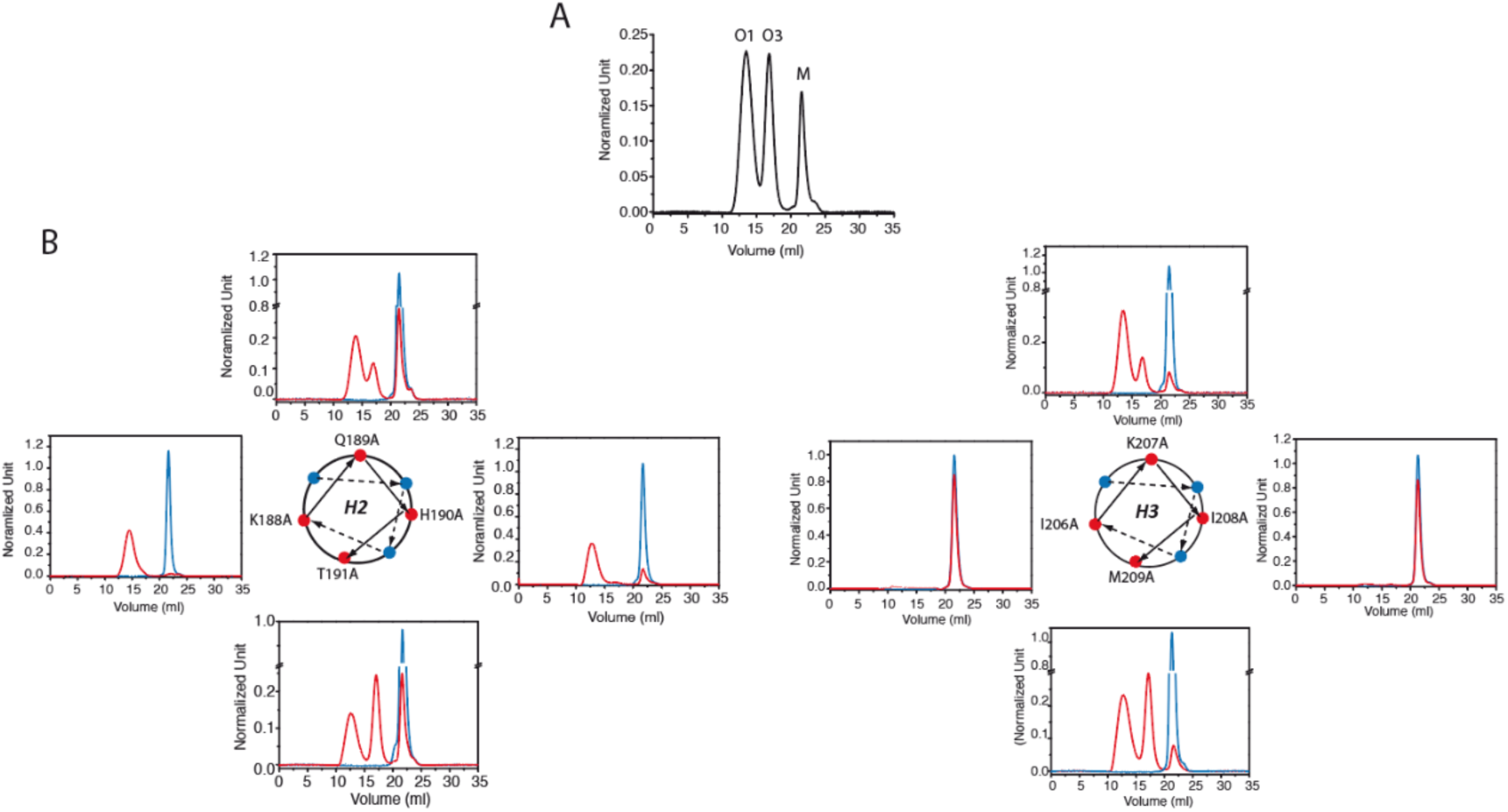
Rational design of an ovine full-length PrP mutant library to modulate oligomerization pathways. (A) Representative size-exclusion chromatography (SEC) profile of wild-type (wt) PrP (100 μM) after incubation at 45 °C for 1 h. Two discrete oligomeric species, O1 (MW∼700kDa) and O3 (MW∼200kDa), are formed through parallel oligomerization pathways, as previously reported (8, 20, 27). The monomeric PrP peak is indicated by M. Single alanine substitutions were introduced within helix H2 (residues 188–191) or helix H3 (residues 206–209) to selectively perturb oligomerization propensity. (B) SEC profiles of individual PrP mutants obtained under identical conditions to those used for wtPrP (incubation at 45°C for 1h). Red traces show oligomerization profiles following incubation, whereas blue traces correspond to the monomeric state of each mutant prior to incubation at 45°C for 1h. Mutations within H2 preferentially bias oligomerization toward the O1 pathway, whereas substitutions within H3 markedly impair PrP self-oligomerization. (C) Conceptual outcomes of heteropolymerization between polymerization-competent PrP variants (blue) and non–self-polymerizing mutants (red). Three scenarios are considered: (1) inhibition of oligomerization through formation of nonproductive heterocomplexes; (2) incorporation of mutant subunits leading to structurally nonequivalent hetero-assemblies; or (3) faithful structural complementation resulting in hetero-assemblies quasi-equivalent to wt homo-assemblies.

To distinguish between these possibilities, wild-type PrP was incubated with increasing concentrations of the non-self-polymerizing mutants I206A or I208A (illustrated for I206A in Fig. 2A). Oligomer formation was monitored by time-resolved size-exclusion chromatography (SEC). In both cases, hetero-oligomerization led to a pronounced increase in the formation of O1 and O3 oligomers, indicating that these mutants are efficiently incorporated into oligomeric assemblies rather than acting as dominant-negative inhibitors. Quantitative mass spectrometry of SEC-purified oligomers further confirmed the concentration-dependent incorporation of I206A and I208A into both O1 and O3 species (Fig.2B and C)

**Figure 2.**
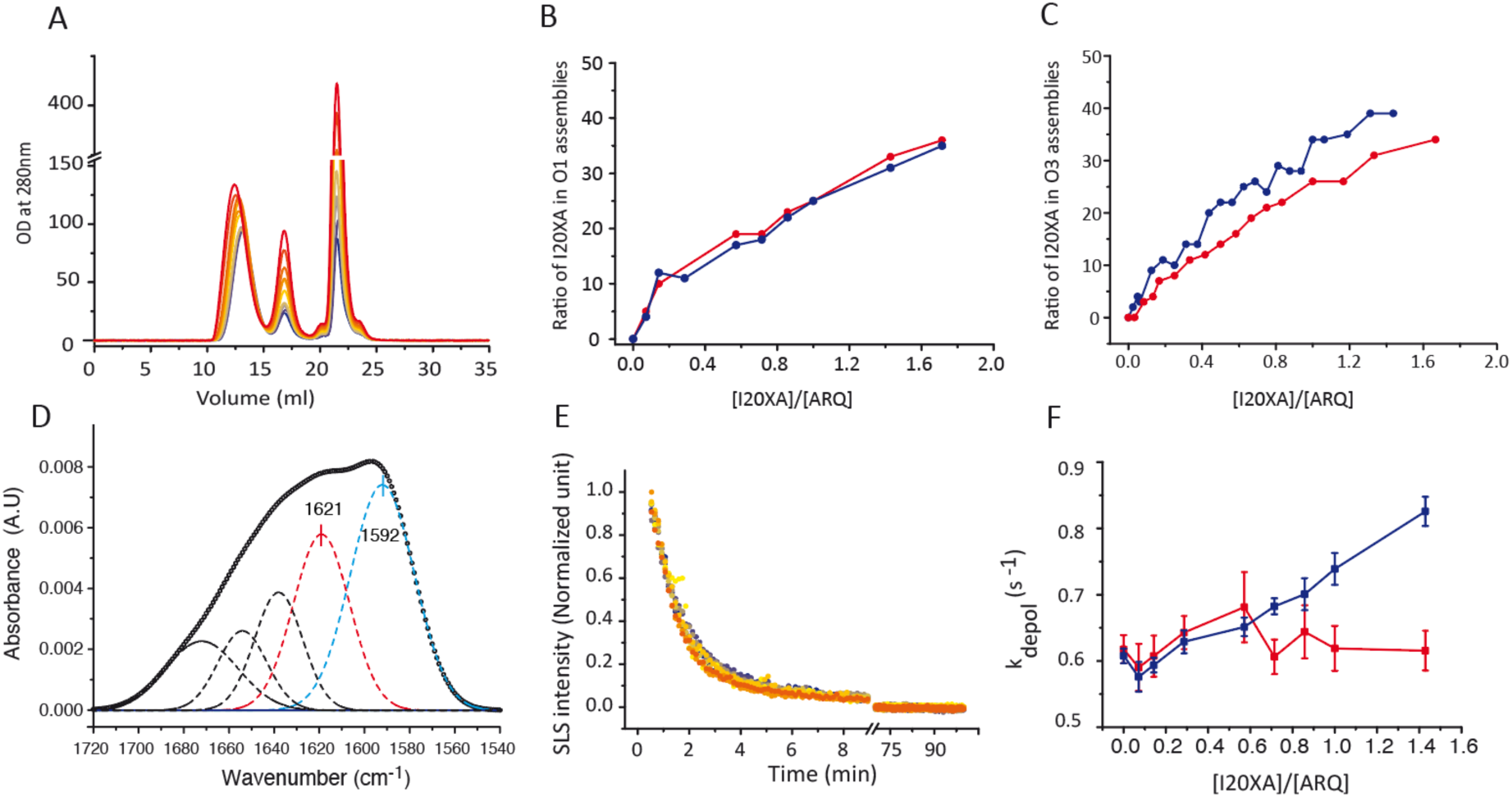
Structural complementation and differential incorporation of non-self-polymerizing PrP mutants into wtPrP oligomers. (A) Typical SEC profiles of wtPrP oligomerization in the presence of increasing concentrations of the non-self-polymerizing mutant I206A. The colour scale indicates I206A concentration, ranging from 0 μM (blue) to 120 μM (red). Increasing I206A concentrations promote the formation of both O1 and O3 oligomers, indicating efficient incorporation into wtPrP assemblies. (B and C) Quantification of mutant incorporation into purified O1 (B) and O3 (C) oligomers. Following SEC purification, the relative amounts of I206A (blue) and I208A (red) integrated into each oligomeric species were determined by LC-FTMS. (D) Secondary-structure analysis of hetero-O1 oligomers composed of uniformly ^13^C-labeled wtPrP and uniformly ^12^C-labeled I206A. FTIR spectra of the amide I region reveal distinct β-sheet-associated bands corresponding to wt and mutant components, indicating that I206A undergoes α-fold to β-fold conformational conversion upon incorporation into O1 assemblies. (E) Depolymerization kinetics of O1 oligomers formed in the presence of increasing concentrations of non-self-polymerizing mutants, monitored by SLS intensity as function of time. (F) Depolymerization rate constants (k_depol_) extracted from SLS measurements for hetero-O1 assemblies containing I206A (blue) or I208A (red). Whereas incorporation of I208A yields hetero-assemblies with stability comparable to wt homo-assemblies, increasing incorporation of I206A results in progressively destabilized assemblies, indicating nonequivalent structural integration.

To determine whether incorporation of non-self-polymerizing mutants is accompanied by acquisition of a β-enriched fold associated with polymerization competence, we performed hetero-oligomerization between uniformly ^13^C-labeled wt PrP and uniformly ^12^C-labeled I206A and subsequently purified the resulting O1 oligomers. These hetero-O1 assemblies were then analysed by FTIR spectroscopy (Fig2.D). Isotopic resolution of the amide I band revealed that both components adopt β-sheet–rich secondary structure within the hetero-assemblies, demonstrating that the mutant undergoes conformational conversion into a polymerization-competent fold upon incorporation.

We next assessed whether hetero-oligomer formation preserves the structural integrity of the oligomeric assemblies by monitoring depolymerization kinetics using static light scattering (Fig.2E). These measurements revealed that incorporation of I208A yields hetero-O1 assemblies with stability comparable to wild-type homo-oligomers, whereas increasing incorporation of I206A resulted in progressively destabilized assemblies (Fig.2F). These distinct outcomes indicate that structural complementation during hetero-oligomerization can be faithful or quasi-faithful, depending on the mutant, and demonstrate that folding information supplied by wild-type PrP is required for mutant incorporation into oligomeric assemblies.

Together, these results demonstrate that during hetero-oligomerization, wild-type PrP transmits folding information to polymerization-defective variants, thereby restoring their ability to adopt a β-enriched, polymerization-competent conformation and to be incorporated into oligomeric assemblies.

### Domain-resolved architecture and non–self-polymerizing mutant segregation in O1 oligomers

Having established that folding information supplied by wt PrP enables incorporation of non-self-polymerizing mutants during oligomerization, we next asked how this information is structurally encoded within the O1 oligomer and whether non-polymerizing mutant incorporation alters its organization and higher-order assembly properties. To address this, O1 oligomers were purified by size-exclusion chromatography and analysed by liquid atomic force microscopy (SI-1). As shown in Fig. 3A, wild type O1 oligomers exhibit pronounced size polydispersity. Atomic force microscopy based single particle analysis revealed that O1 oligomers adopt a two-domain architecture composed of a large globular domain of approximately 8nm in diameter, hereafter referred to as the B domain, connected to an elongated domain, the E domain, approximately 3nm in diameter and of variable length (Fig. 3B and C). The E domain was observed to associate with either face of the B domain, suggesting a quasi-equivalence of B domain surfaces with respect to E domain attachment. Additional configurations were detected, including isolated E domains and repetitive modular arrangements composed of successive B and E domains (Fig. 3C).

**Figure 3.**
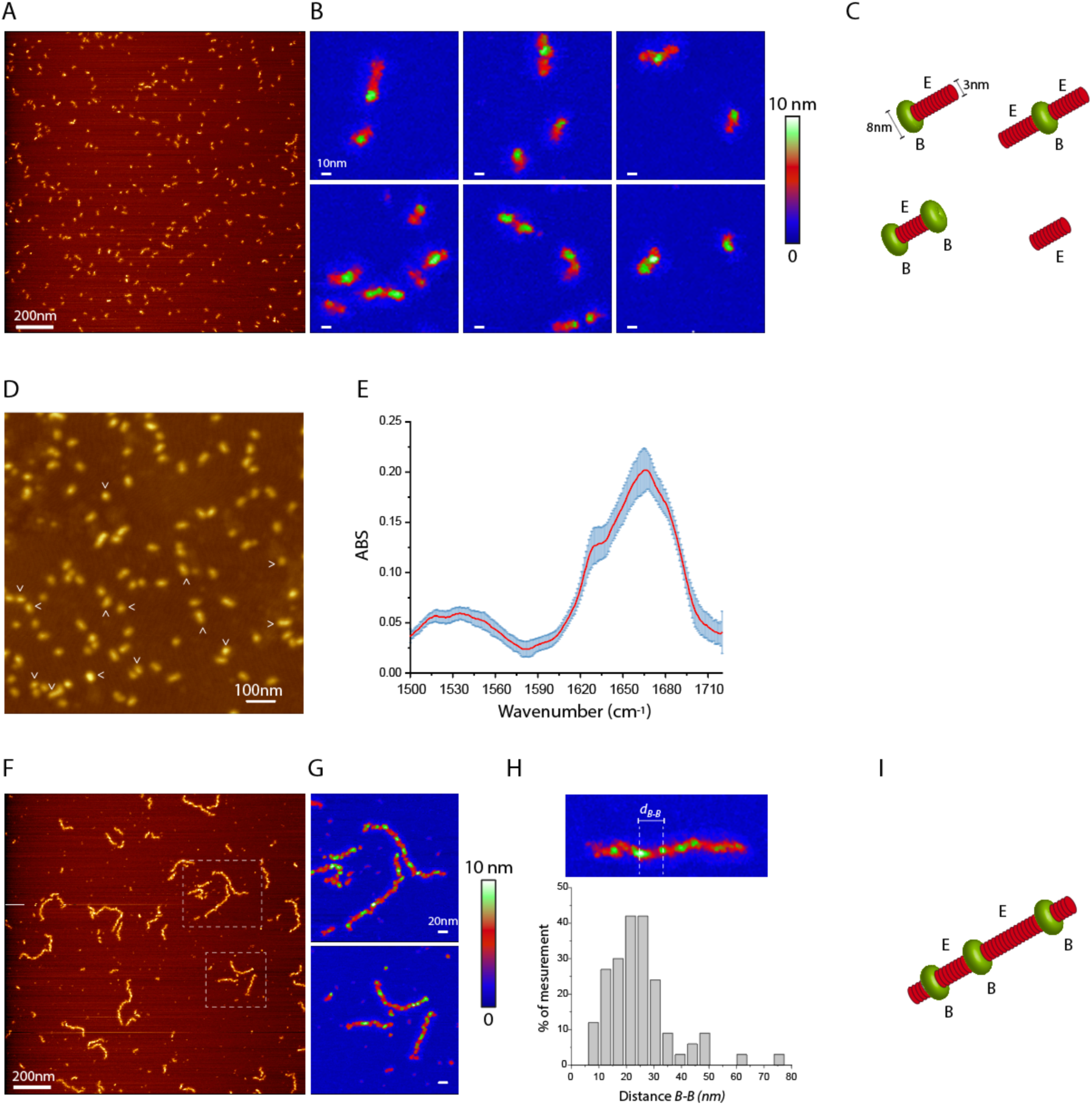
Domain-resolved topology of wt O1 oligomers revealed by high-resolution AFM. (A) High-resolution liquid AFM images of wild-type (wt) O1 oligomers following purification by size-exclusion chromatography, illustrating the pronounced size and shape heterogeneity of individual particles. (B) Representative classes of O1 oligomer topologies identified by single-particle AFM analysis. O1 assemblies display a modular organization composed of a compact globular domain (B domain) associated with one or more elongated domains (E domains), as well as isolated or repetitive modular configurations. (C) Schematic topological models corresponding to each experimentally observed O1 oligomer subclass shown in (B), highlighting the relative organization and connectivity of B and E domains within the oligomeric architecture. (D) AFM topographic image of wt O1 oligomers used to guide single-particle AFM-IR measurements. Arrows indicate representative individual O1 assemblies selected for infrared spectral acquisition on a particle-by-particle basis. (E) Mean infrared absorption spectra in the amide I and amide II’ regions recorded from the individual O1 oligomers indicated in (D). The Spectrum represent the average of measurements acquired from 15 single particles, with shaded areas indicating the corresponding standard deviation. The presence of a shoulder centred at ∼1629 cm⁻¹ is indicative of β-strand-rich structural elements within the oligomeric assemblies. (F) High-resolution liquid AFM image of the condensed state of wt O1 oligomers induced by ultracentrifugation (see Materials and Methods and SI-2). Under these conditions, O1 oligomers undergo higher-order association to form elongated, worm-like supramolecular assemblies. (G) Topological analysis of condensed O1 assemblies reveals an aperiodic succession of globular B domains and elongated E domains, consistent with a modular condensation process characterized by repeated – B–E– organization along the assembly axis. (H) Distribution of center-to-center distances between adjacent B domains within condensed O1 assemblies, revealing a nonperiodic spacing consistent with the aperiodic-B-E-modular organization observed in panels F and G. (I) Schematic topological model of the condensed O1 assemblies illustrating the nonperiodic, linear concatenation of B and E domains inferred from AFM analysis.

The secondary structure of O1 oligomers was further examined at the level of individual oligomeric assemblies using single-particle AFM-IR spectroscopy (Fig. 3D and E). Infrared spectra acquired from selected single assemblies consistently revealed the presence of a shoulder feature centred at ∼1629 cm⁻¹ in the amide I region, indicative of β-sheet-rich structural elements within O1 oligomers. Owing to the effective spatial resolution imposed by the relatively large radius of the AFM-IR cantilever, it was not possible to resolve the secondary structure of the B and E domains independently within individual assemblies. Consequently, the observed β-sheet signature reflects the average secondary structure of whole O1 oligomers rather than domain-specific contributions. The measurable standard deviation (Fig. 3E) associated with the averaged spectra may nevertheless reflect underlying secondary-structure heterogeneity within O1 assemblies, consistent with the coexistence of structurally distinct B and E domains. This modular two domain organization is consistent with our previously proposed structural model of the O1 oligomer (24). Upon increasing local concentration by ultracentrifugation (see SI-2), wild-type O1 oligomers underwent modular condensation to form elongated, worm-like, non-anastomosed fibrillar assemblies. High-resolution AFM analysis revealed that these supramolecular structures are composed of an aperiodic succession of compact globular B domains interconnected by elongated E domains, resulting in a linear but nonperiodic organization of-E-B-E-modules along the assembly axis (Fig. 3F to H).

To determine at the single molecule level whether the non-self-polymerizing mutant is incorporated randomly or preferentially into a specific O1 domain, we employed an atomic force microscopy infrared approach combined with isotopic labelling. Wt PrP was uniformly labelled with ^13^C, whereas the non-polymerizing I206A mutant retained the natural ^12^C isotope. This isotopic differentiation enabled discrimination between the amide one infrared signatures of wild type and mutant PrP, thereby allowing spatial localization of each species within individual hetero oligomers (see Materials and Methods).

During heteropolymerization, ^13^C labelled wild type PrP and ^12^C I206A formed hetero O1 oligomers with incorporation efficiencies comparable to those observed in non-isotopically labelled systems and retained the ability to undergo ultracentrifugation induced condensation into worm like fibrillar assemblies (Fig. 4A and B). Both the morphology of hetero O1 oligomers and that of their condensed fibrillar assemblies were indistinguishable from the corresponding wild type homo assemblies (Fig. 4C and Fig. 3).

**Figure 4.**
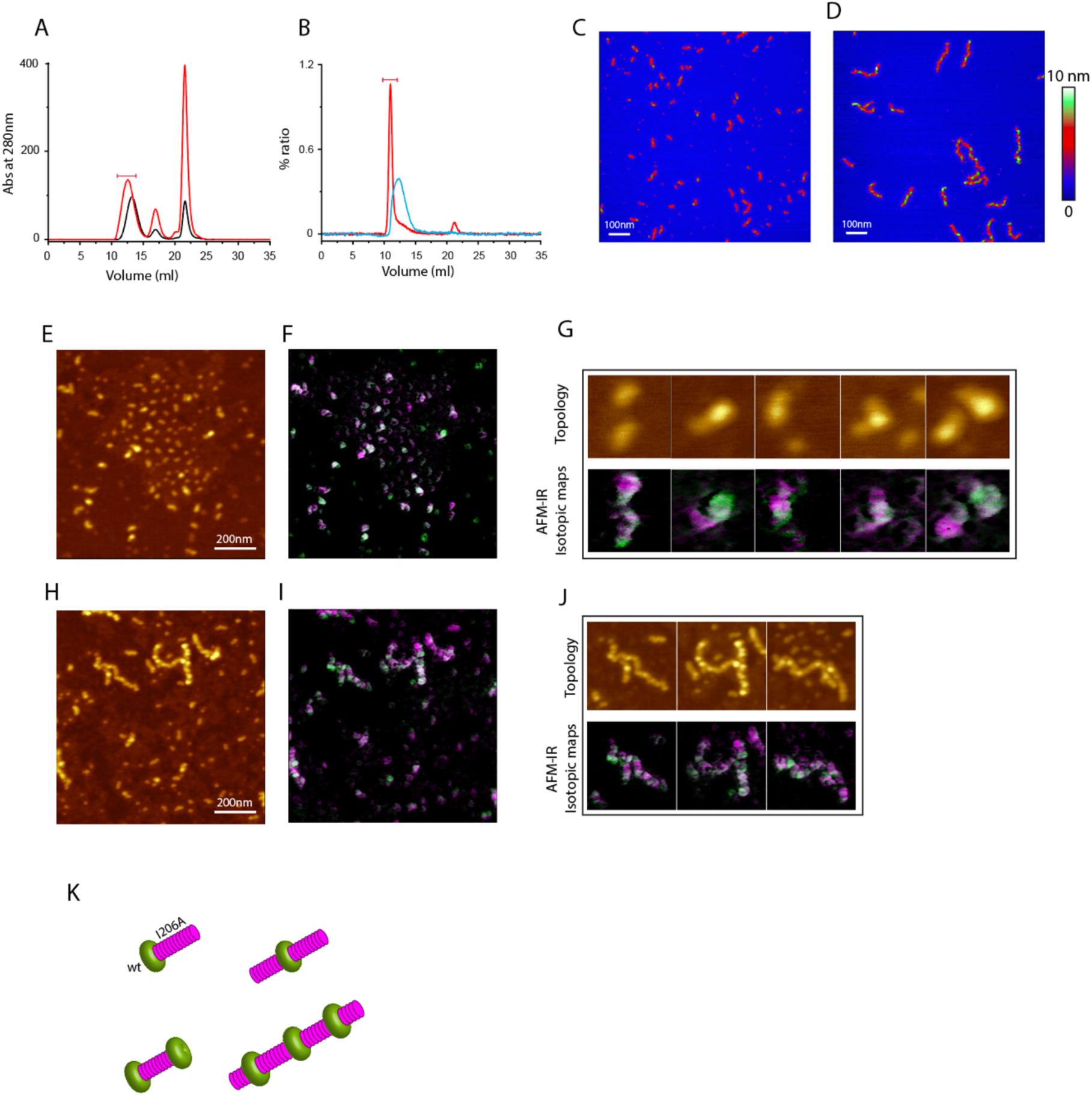
Domain-specific partitioning of wtPrP and I206A within hetero-O1 oligomers and their condensed assemblies revealed by AFM-IR. (A) SEC profiles of uniformly ^13^C-labeled wild-type PrP (in black line) and ^13^C-wtPrP copolymerized with one molar equivalent of uniformly ^12^C-labeled I206A (in red line). The red bar above the O1 peak indicates the SEC fraction collected for subsequent analyses. (B) SEC profiles of purified hetero-O1 oligomers before (in blue line) and after (in red line) ultracentrifugation, which induces condensation of O1 assemblies. The red bar indicates the fraction collected for downstream AFM and AFM-IR analyses. (C and D) High-resolution liquid AFM images of hetero-O1 oligomers prior to condensation (C) and of condensed hetero-O1 assemblies following ultracentrifugation (D), showing worm-like supramolecular architectures. (E and F) AFM topography (E) and corresponding AFM-IR hyperspectral maps (F) of hetero-O1 oligomers, highlighting the spatial distribution of amide I absorption bands characteristic of ^13^C-labeled wtPrP (green) and ^12^C-labeled I206A (pink). (G) Representative AFM topography and corresponding AFM-IR spectral maps of selected hetero-O1 assemblies, illustrating domain-specific localization of ^13^C-wtPrP and ^12^C-I206A within individual oligomers. (H and I) AFM topography (H) and corresponding AFM-IR hyperspectral maps (I) of condensed hetero-O1 assemblies, revealing preservation of isotopic segregation following condensation. (J) Representative AFM topography and AFM-IR spectral maps of selected condensed hetero-O1 assemblies, highlighting the spatial partitioning of ^13^C-wtPrP and ^12^C-I206A along worm-like architectures. (K) Schematic representation summarizing the domain-biased partitioning of wild-type PrP and the non-self-polymerizing I206A mutant within hetero-O1 oligomers and their condensed assemblies. Wild-type PrP predominantly localizes within the globular B domain. The I206A mutant preferentially partitions into the elongated E domain, while remaining permissive for incorporation within, or in close proximity to, the B domain in a subset of assemblies, particularly highlighted in the condensate state of hetero O1. Together, these observations support a two-step assembly mechanism in which folding information transferred from wild-type PrP enables structural recruitment of non-self-polymerizing variants, followed by domain-biased organization within the mature oligomeric architecture.

Hyperspectral infrared imaging of hetero-O1 oligomers (Fig. 4E–G) and their worm-like condensates (Fig. 4H–J) revealed a non-random spatial organization of the two isotopically labeled species (SI-3). Using reference amide I absorption bands centered at approximately 1590 cm⁻¹ for uniformly ¹³C-labeled wild-type PrP and at approximately 1690 cm⁻¹ for ¹²C-labeled I206A, we observed pronounced spatial segregation between wild-type and mutant protomers within individual assemblies (Fig. 4K).Importantly, while I206A was predominantly localized within the elongated E domain, AFM-IR mapping also revealed its presence within, or in close proximity to, the globular B domain in a subset of hetero-O1 assemblies, particularly in the condensed state (Fig. 4J). These observations indicate that incorporation of the non-self-polymerizing mutant does not occur through simple passive partitioning, but rather involves folding information transfer from wild-type protomers that enables I206A to adopt conformations compatible with integration into both structural domains of the oligomer.

Together, these data support a two-step mechanism in which I206A is first structurally recruited during hetero-oligomerization through direct interaction with wild-type PrP, followed by domain-specific organization within the mature oligomeric architecture. This organization is biased toward segregation within the E domain, while remaining permissive for incorporation within the B domain under conditions of higher local concentration and oligomer condensation.

### Transient Conformers Enable Polymerization of Inactive Mutants under arrested reaction condition

The formation of hetero-assemblies through incorporation of non–self-polymerizing mutants such as I206A or I208A could occur via transient folding intermediates that compensate for the missing polymerization-competent folding information of these mutants. To test this hypothesis, heteropolymerization assays were performed under conditions in which wild-type PrP alone does not form detectable oligomers within the experimental timescale. This arrested reaction condition (ARC) regime exploits a general kinetic property of polymerization reactions, whereby reaction orders greater than one render oligomer formation negligible at sufficiently low reactant concentrations, despite the possible transient formation of short-lived monomeric or oligomeric intermediates (Fig. 5A).

**Figure 5.**
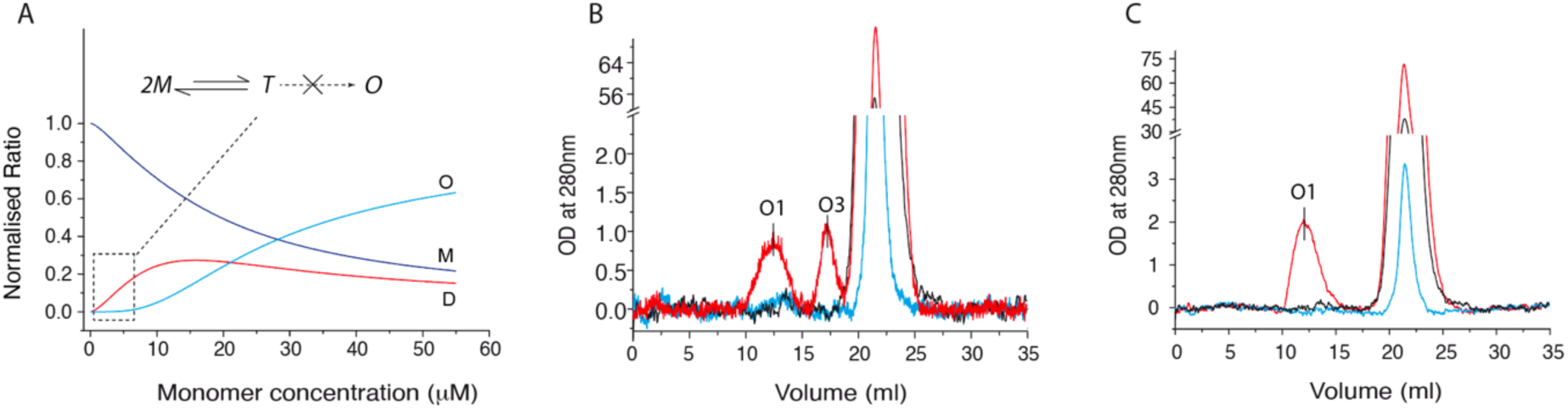
Transient conformers enable polymerization of non-self-polymerizing mutants under arrested reaction conditions (ARC). (A) Representative simulation of a two-step, second-order oligomerization reaction illustrating the ARC regime. At sufficiently low initial monomer concentrations, formation of stable oligomeric species (O) is kinetically suppressed, whereas transient intermediate species (T) can be transiently populated. The relative abundance of each species is plotted as a function of initial monomer concentration. (B) SEC profiles following incubation at 50 °C of wild-type (wt) PrP at subcritical concentration (3μM) and the non–self-polymerizing mutant I206A (50μM). Neither wtPrP alone (black trace) nor I206A alone (blue trace) forms detectable oligomers under these conditions, whereas co-incubation of wtPrP and I206A (red trace) restores formation of both O1 and O3 oligomers. (C) SEC profiles following incubation at 50 °C of the selective oligomerizing mutant H190A (3μM) and I206A (50μM). H190A alone (black trace) and I206A alone (blue trace) remain monomeric, whereas co-incubation (red trace) selectively yields O1 oligomers, consistent with pathway-specific structural information transfer under ARC conditions.

For wild-type PrP, ARC correspond to monomer concentrations ≤3 μM, under which incubation at 50°C does not lead to detectable formation of either O1 or O3 oligomers, as assessed by SEC (Fig. 5B, black trace). Likewise, the non-self-polymerizing mutant I206A remains strictly monomeric when incubated alone at high concentration under the same conditions (Fig. 5B, blue trace). Strikingly, co-incubation of subcritical concentrations of wtPrP with excess I206A resulted in the appearance of both O1 and O3 oligomeric species (Fig. 5B, red trace). Because neither component polymerizes independently under ARC conditions, this restoration of oligomer formation, even at low abundance, indicates that wtPrP transiently populates a polymerization-competent conformational state capable of transferring folding information to I206A, thereby enabling oligomer assembly.

A similar behaviour was observed when wtPrP was replaced by selective oligomerizing mutants. When the H190A mutant, which oligomerized exclusively through the O1 pathway (Fig. 1B), was incubated under ARC conditions with I206A at 50 μM, only O1 oligomers were detected. Notably, neither H190A at 3μM nor I206A at 50μM polymerized when incubated alone (Fig. 5C, red trace). Comparable results were obtained when the K188A mutant was used in place of H190A (SI-3).

Together, these results tend to suggest that polymerization-competent PrP variants, including wtPrP and selective oligomerizing mutants such as H190A or K188A, can generate transient conformers capable of transferring folding information to non-self-polymerizing mutants. This complementation restores the missing folding information required for oligomerization and initiates assembly under conditions where neither component polymerizes independently.

### O1 Oligomer restores the polymerization of I206A by templating

Having established that non–self-polymerizing mutants can be incorporated into oligomeric assemblies through structural complementation during oligomerization, we next asked whether this process can also occur through a templating mechanism mediated by preformed oligomers. In this context, we hypothesized that O1 oligomers, and more specifically defined structural domains within their modular architecture (B and E), could function as conformational templates that promote folding of the non-self-polymerizing I206A mutant through direct interaction, thereby enabling its incorporation into hetero-O1 assemblies.

To assess the integration of the I206A mutant into O1 oligomers and to precisely localize its incorporation with respect to the B and E domains of O1, an AFM-IR approach was employed. Uniformly ^12^C-labeled I206A was incubated with uniformly ^13^C-labeled O1 oligomers at 50 °C. Changes in the mean weight-average molecular mass of the assemblies were monitored by measuring static light-scattering intensity (Fig. 6A). As shown, the scattering intensity of the ^13^C-O1/^12^C-I206A mixture increased nonlinearly, following a power-law relationship, whereas the scattering intensities of ^12^C-I206A or ^13^C-O1 alone did not vary significantly over the same time scale. The products of the ^13^C-O1/^12^C-I206A incubation were subsequently passed through a size-exclusion chromatography (SEC) column to purify large assemblies and separate them from unreacted ^12^C-I206A (see also SI-1) prior to AFM-IR analysis. Following purification, hyperspectral mapping revealed clear colocalization of the 1650cm⁻¹ signal with the 1590cm⁻¹ absorption band within the same oligomers, indicating incorporation of the I206A mutant into wild-type O1 assemblies (Fig. 6B and C). Spectral analysis at the level of individual assemblies allowed the identification of at least three distinct organizational categories (Fig.6D), depending on whether I206A was incorporated in the vicinity of the B or E domains within the wtO1 assemblies. Notably, a small number of assemblies lacking the 1590 cm⁻¹ absorption band were also observed, indicating the formation of oligomeric assemblies devoid of I206A incorporation.

**Figure 6.**
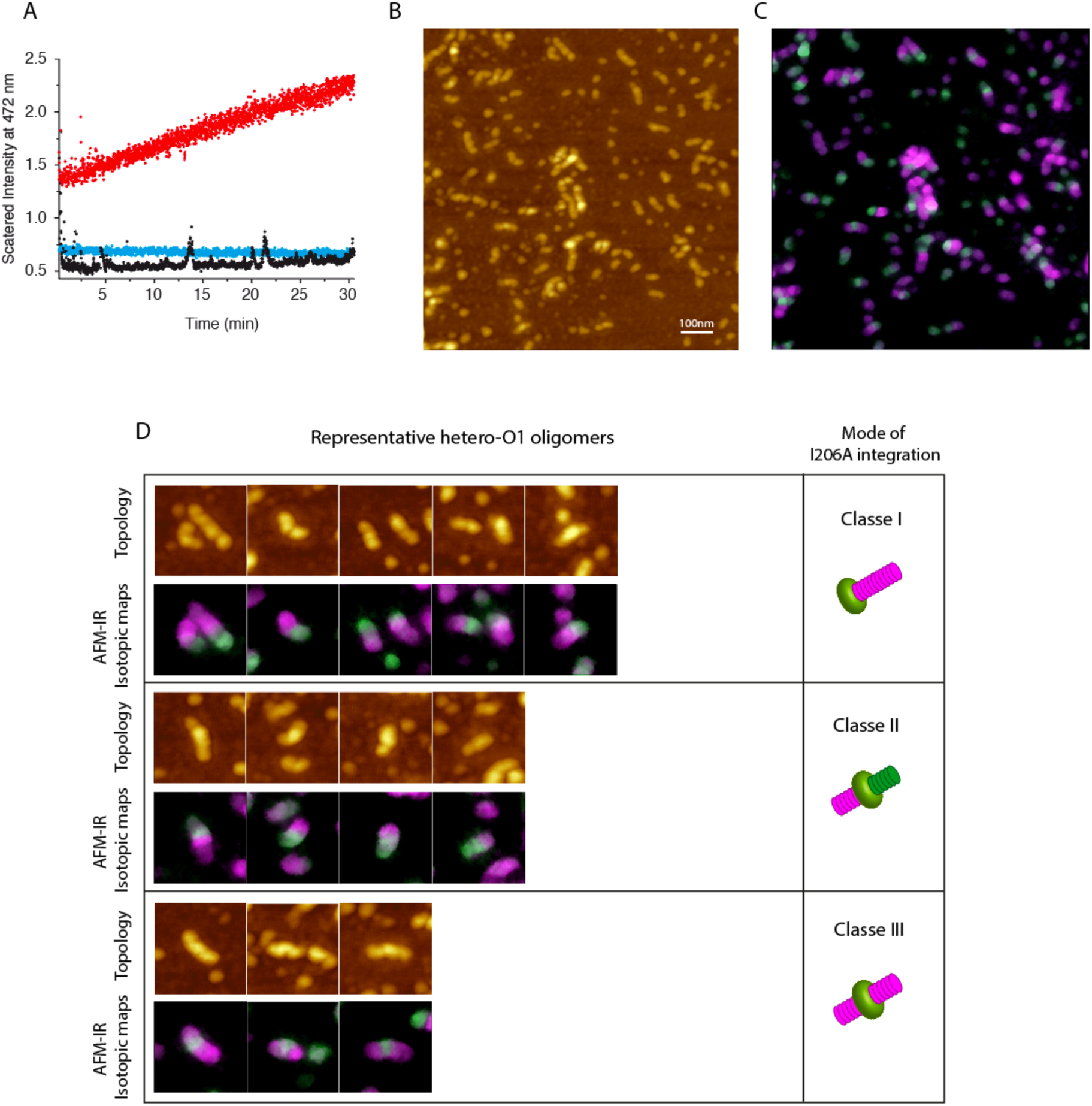
Preformed O1 oligomers template incorporation of the non–self-polymerizing I206A mutant. (A) Static light-scattering (SLS) intensity measured as a function of time at 42 °C for uniformly ^12^C-labeled I206A (30 μM; black trace), purified uniformly ^13^C-labeled O1 oligomers (3 μM, expressed as monomer equivalents; blue trace), and a mixture of ^13^C-labeled O1 (3 μM) with ^12^C-labeled I206A (red trace). Whereas neither O1 nor I206A alone exhibits a significant change in scattering intensity over time, the O1/I206A mixture displays a progressive increase in scattering intensity, indicating an increase in the weight-average molecular mass of assemblies formed in solution. (B and C) AFM topography (B) and corresponding AFM-IR hyperspectral maps (C) of SEC-purified assemblies obtained after incubation of ^13^C-labeled O1 with ^12^C-labeled I206A at 42 °C. AFM-IR maps recorded at 1590 cm⁻¹ and 1690 cm⁻¹ selectively localize the amide I absorption of ^13^C-labeled wtPrP (in green) and ^12^C-labeled I206A (in pink), respectively, revealing spatial partitioning of the two species within hetero-O1 assemblies. (D) Single-particle AFM-IR analysis of SEC-purified products formed upon incubation of ^13^C-labeled O1 oligomers with ^12^C-labeled I206A reveals three distinct modes of mutant incorporation. For each mode, representative AFM topography images and corresponding AFM-IR isotopic maps show the spatial distributions of wtPrP (^13^C, green) and I206A (^12^C, pink) within individual hetero-O1 assemblies. These modes correspond to preferential localization of I206A in the vicinity of the β-sheet-rich B domain, at the B/E domain interface, or within the elongated E domain, indicating distinct patterns of structural complementation by the O1 scaffold. Together, these single-particle data show that incorporation of the non–self-polymerizing mutant is governed by domain-specific structural information transfer from wtPrP, rather than by random association.

Taken together, these results demonstrate that O1 oligomers act as conformational templates for the non-self-polymerizing I206A mutant, transmitting the folding information required for its conversion into a polymerization-competent state. AFM-IR isotopic mapping indicates that this templating activity is primarily mediated by B domain, which serves as the principal initiation scaffold for folding information transfer and mutant incorporation. Nevertheless, owing to the intrinsic spatial resolution limits of AFM-IR, we cannot exclude the existence of a minor subpopulation in which I206A is also associated with the elongated E domain of wtO1 assemblies. Such configurations may correspond to secondary recruitment events following initial B-domain-mediated templating or to alternative modes of domain engagement that remain below the current resolution threshold.

## DISCUSSION

The prion phenomenon relies on the autonomous propagation of structural information through one or multiple protein conformational transitions. It is generally accepted that this propagation predominantly occurs via fibril end-elongation. However, the capacity of alternative quaternary fold such as oligomers or transient folding intermediates to initiate or propagate conformational information remains poorly understood. The generation of PrP single mutant in which self-oligomerization is selectively suppressed or modulated (20) enables investigation of mechanisms that extend beyond simple aggregation driven by protein misfolding. This approach allows exploration of folding information transfer and complementation mediated by non-fibrillar assemblies, including both stable oligomeric species and transient folding intermediates.

AFM imaging of O1 oligomers reveals a modular architecture composed of two distinct domains, termed the B and E domains (Fig. 3C), with the B domain forming a compact structural core. Previous studies demonstrated that formation of both O1 and O3 oligomers strictly requires conformational rearrangements within the H2H3 region of PrP, whereas deletion of the S1H1S2 segment does not impair oligomer assembly (8, 19–21, 24). These observations effectively restrict the oligomerization-competent region of PrP to H2H3 and suggest that the B domain corresponds to the minimal structural core driving oligomer formation. In this context, molecular dynamics simulations focused on the isolated H2H3 domain have shown that PrP oligomerization proceeds through a noncanonical two-step mechanism, in which a stable oligomeric “basement” of approximately 5–8 monomers form first and acts as an initiation core for subsequent growth (24). We propose that the B domain identified experimentally represents the structural counterpart of this oligomeric basement, whereas the E domain reflects later-stage accretion of additional PrP units as linear or branched extensions (Fig.7). While this correspondence remains inferential and awaits direct structural validation at atomic resolution, the convergence between AFM topology and MD-derived assembly mechanisms provides a coherent framework for interpreting early PrP oligomerization events.

**Figure 7.**
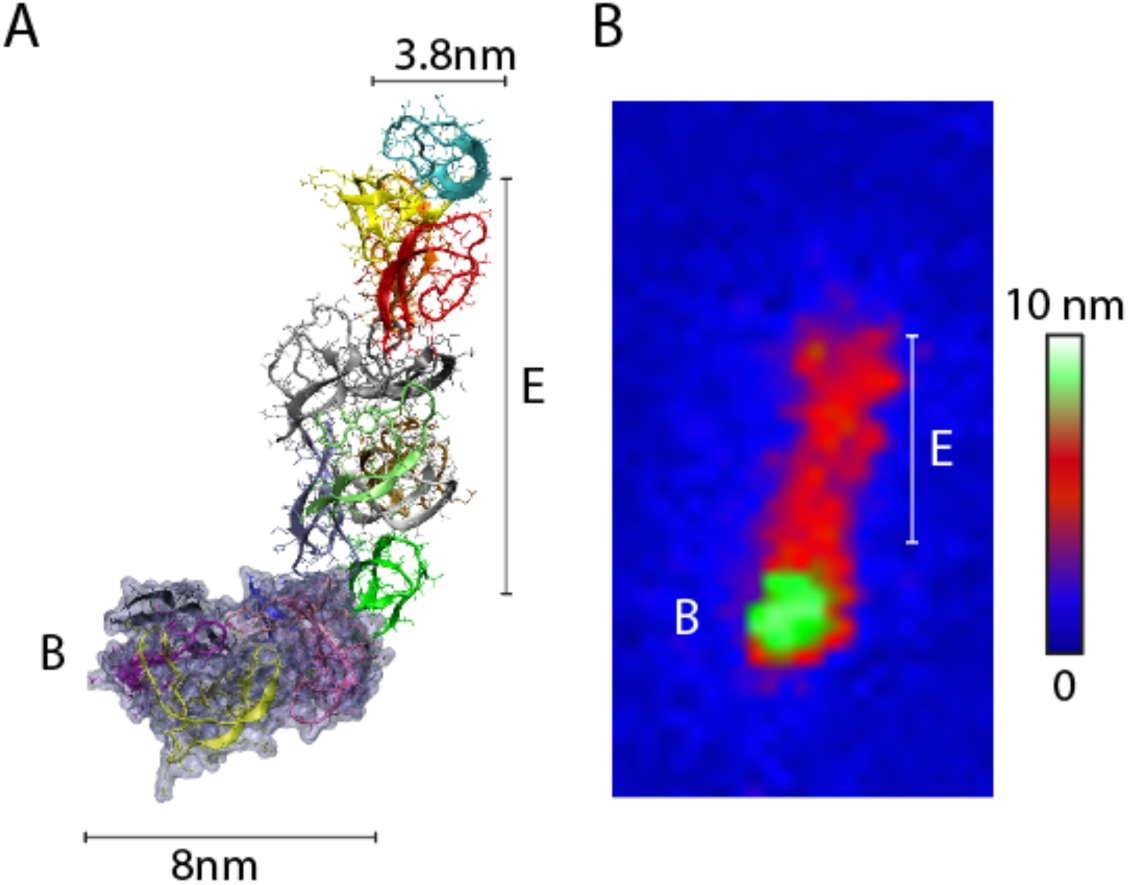
Convergence between molecular dynamics–derived oligomerization models and experimentally observed O1 topology. (A) Schematic representation of the two-step oligomerization mechanism proposed by Collu et al (24). based on atomistic molecular dynamics simulations of the β-rich H2H3 domain of ovine PrP. In this model, oligomerization proceeds through the formation of a compact, stable oligomeric “basement” composed of approximately 5-8 PrP units, which acts as an initiation core. Subsequent growth occurs via the attachment of additional PrP units or small oligomers as quasi-linear extensions (“branches”), giving rise to larger, non-fibrillar assemblies. (B) High-resolution liquid AFM image of purified O1 oligomers highlighting a modular two-domain organization characterized by a compact globular B domain associated with elongated E domains. This experimentally observed topology is compatible with the basement-and-extension architecture described by Collu et al. (24), supporting the correspondence between the B domain identified here and the oligomeric basement predicted by molecular dynamics simulations. Together, these observations support a model in which the β-sheet-rich B domain of O1 assemblies constitutes a functional oligomeric basement that initiates growth and organizes subsequent domain accretion.

During heteropolymerization experiments, non–self-polymerizing PrP variants are efficiently incorporated into oligomeric assemblies and undergo a conformational transition from an α-helix–enriched fold to a β-sheet–rich architecture, demonstrating direct transfer of folding information from wtPrP to otherwise inactive mutants. Single-particle AFM-IR analyses further reveal that protomer organization within hetero-O1 assemblies is non-random and reflects domain-level structural organization. While the I206A mutant exhibits a pronounced tendency to segregate from wtPrP and preferentially localizes within the elongated E domain, AFM-IR mapping also reveals its presence within, or in close proximity to, the globular B domain in a subset of assemblies, particularly following oligomer condensation (Fig. 4J, K). This observation indicates that folding information supplied by wtPrP during hetero-oligomerization can enable I206A to adopt conformations compatible with integration into both structural domains of the O1 architecture. These findings support a two-step mechanism in which mutant protomers are first structurally recruited through folding information transfer from wtPrP, followed by domain-biased spatial organization within the mature oligomer. In this framework, the B domain functions as the primary initiation and information-donating scaffold, whereas the E domain constitutes a more permissive structural environment that preferentially accommodates mutant incorporation, particularly under conditions of increased local concentration and oligomer condensation.

Taken together with the oligomerization mechanism inferred from molecular dynamics simulations (24), these observations provide a structural rationale for the intrinsic inability of non-self-polymerizing mutants such as I206A and I208A to oligomerize autonomously. One plausible explanation for the inability of the I206A mutant to oligomerize autonomously is its failure to nucleate alone the formation of the initial oligomeric basement, corresponding to the β-sheet-rich B domain that functions as the initiation core of O1 assemblies. Under heteropolymerization conditions, wtPrP overcomes this limitation by forming the B domain scaffold, which subsequently recruits mutant protomers and promotes their conformational conversion and polymerization.

The arrested reaction condition further indicates that polymerization-competent PrP variants, such as wtPrP and H190A, can transiently adopt conformations that compensate for the missing folding information of the non-self-polymerizing I206A mutant and thereby enable oligomer formation. These transient conformers likely correspond to unstable oligomeric states whose lifetime and accumulation critically depend on the initial monomer concentration. Consistent with this interpretation, wtPrP, capable of accessing both the O1 and O3 oligomerization pathways, promotes the formation of both O1 and O3 assemblies in the presence of I206A under ARC conditions, whereas the H190A mutant, whose oligomerization landscape is restricted to the O1 pathway, selectively supports formation of O1 assemblies. These observations indicate that the transient conformers populated by wtPrP and H190A encode distinct, pathway-specific folding information that can be transmitted to the I206A mutant, thereby directing the nature of the oligomeric assemblies that emerge.

The complementation process and transfer of structural information is further substantiated when preformed wtO1 oligomers are incubated with the non-self-polymerizing I206A mutant. As shown in Fig.6D, single-particle AFM-IR analysis reveals a clear spatial partitioning between ^13^C-labeled wt O1 and ^12^C-labeled I206A within individual hetero-assemblies. Although the spatial resolution of AFM-IR is intrinsically lower than that of high-resolution liquid AFM (primarily due to the larger effective radius of cantilever used for AFM-IR) correlative analysis combining topographical features with local infrared spectra consistently indicates that I206A incorporation occurs preferentially in the vicinity of the B domain. These observations indicate that the B domain of the O1 oligomer constitutes a functional initiation interface capable of restoring polymerization competence to an otherwise inactive PrP variant. By promoting the conformational conversion of I206A through a templated, induced-fit mechanism, the B domain acts as a scaffold for structural information transfer.

## Conclusions

The prion paradigm has long emphasized fibril ends as the exclusive sites for conformational templating and polymerization. The present study expands this view by demonstrating that non-fibrillar quaternary assemblies can also store and transmit folding information at early stages of PrP oligomerization. Using recombinant PrP under controlled *in vitro* conditions, we show that discrete O1 oligomers act as active conformational templates capable of restoring polymerization competence to otherwise inactive mutants. Structural and isotopic analyses identify a B domain as the principal initiation scaffold for this process, consistent with previous descriptions of stable early oligomeric cores during prion assembly. We further show that O1 oligomers can undergo modular condensation into higher-order assemblies through accretion of an elongated, structurally distinct E domain, supporting a hierarchical assembly pathway in which folding information transmitted by the B domain initiates subsequent structural elaboration. In this framework, the B domain may represent an oligomer-based secondary nucleation platform, distinct from classical fibril ends, that concentrates and conformationally primes PrP monomers for further assembly.

The marked structural divergence between B and E domains further suggests that early oligomeric intermediates may constitute privileged sites for spontaneous quaternary diversification, potentially contributing to the emergence of alternative assembly pathways. While our conclusions are necessarily limited by the use of recombinant PrP and oligomers generated under defined in-vitro conditions, they establish a proof-of-principle that fibril ends are not the sole initiators of prion polymerization. More broadly, these findings support the idea that early, non-fibrillar assemblies can play an active role in initiating, diversifying, and propagating misfolded protein states in prion diseases and related proteinopathies.

## MATERIAL AND METHODS

### PrP production and purification

Full-length ovine prion protein (PrP; A136, R,154, Q171 variant) and point mutants were produced as previously described (25). Proteins were expressed in Escherichia coli BL21 cells, recovered from inclusion bodies, solubilized in 6 M guanidine hydrochloride containing 10 mM imidazole (pH 7.1), and purified by Ni–NTA affinity chromatography (GE Healthcare). Eluted proteins were desalted into ammonium acetate (0.5 g·L⁻¹, pH 4.6), flash-frozen, lyophilized, resuspended in 20 mM sodium acetate (pH 5.0), and further desalted on a G25 column equilibrated with 20 mM sodium citrate (pH 3.5). Protein concentrations were determined by absorbance at 280 nm using a molar extinction coefficient of 59,485 M⁻¹·cm⁻¹. Uniformly ^13^C-labeled wild-type PrP was expressed in E. coli BL21(DE3). A 5-mL LB preculture was used to inoculate M9 minimal medium supplemented with 1 g·L⁻¹ NH₄Cl, 2 g·L⁻¹ uniformly ^13^C-labeled glucose, and ampicillin. Cultures were grown at 37 °C to mid-log phase and protein expression was induced with 100 μM IPTG for 12 h. Labelled proteins were purified using the same protocol as for unlabelled PrP.

### Oligomer and hetero-oligomers purification and isotope labelling

O1 oligomers were prepared as previously described (8). Briefly, wild-type (wt) PrP (80 μM) in 20 mM sodium citrate buffer (pH 3.5) was incubated at 50 °C for 60 min and subsequently fractionated by size-exclusion chromatography (SEC) using a TSK4000SW column equilibrated in the same buffer. Fractions corresponding to O1 oligomers were collected based on their characteristic elution volume (see also SI-1). Oligomer concentrations were estimated by absorbance at 280 nm using a molar extinction coefficient of 59,485 M⁻¹·cm⁻¹ (expressed as monomer equivalents). Purified O1 oligomers were stored at 4 °C and used within 48 h for subsequent analyses. Uniformly ^13^C-labeled O1 oligomers were generated using uniformly ^13^C-labeled wtPrP monomers following the same protocol. Hetero O1 oligomers were prepared by mixing wtPrP with non-self-polymerizing mutants at the indicated molar ratios prior to incubation at 50 °C. Following incubation, hetero-oligomers were purified by SEC using the same conditions as for wt O1 assemblies, and fractions were collected as indicated in the corresponding figure panels. Condensation of O1 oligomers was induced by increasing their local concentration using an ultracentrifugation-based isopycnic concentration approach (8). SEC purified wild-type or hetero-O1 oligomers were subjected to ultracentrifugation in a Beckman TL100.1 rotor for 4 h at 15 °C. Following centrifugation, the entire solution volume was gently homogenized to ensure uniform redistribution of concentrated oligomeric assemblies. The condensed O1 oligomers were subsequently re-purified by SEC prior to downstream analyses (see also SI-2).

### Size exclusion chromatography (SEC) for polymerization profile and oligomer purification

All size-exclusion chromatography (SEC) experiments were performed using a TSK4000SW column (60 cm; Tosoh Bioscience) equilibrated in 20 mM sodium citrate buffer (pH 3.5). Separations were carried out at room temperature at a flow rate of 1 mL·min⁻¹. For both oligomerization profile analysis and oligomer purification, a sample volume of 500 μL was injected using a 500μL sample loop. Protein elution was monitored by absorbance at 280 nm.

### Mass spectrometry

Mass spectrometry experiments were performed on a QTOF Premier (Waters) mass spectrometer with a nano-ESI ion source probe. The purified oligomers were buffer exchanged against 10 mM triethylammonium (pH 3.5) buffer using MicroSpin G-25 columns. After exchange, the solution was diluted 1:1 with Acetonitrile / Formic acid (99:1 v/v) in order to denature the oligomer and its proteins. The sample was then loaded in a Proxeon metal-coated capillary tip which was inserted in the nano-ESI ion source. After acquisition of the mass spectra of the denatured protein, MaxEnt software was used to deconvolve the charge state distribution. The relative abundance of each monomeric specie in the oligomer was determined based on the integration of the peak corresponding to each protein. To take into account the presence of adducts (< 10% of the intensity) of the lowest mass protein interfering with the peak of the higher mass one, a correction was done to the highest mass protein intensity by removing the contribution of the lower mass one considering the relative abundance of adducts observed in a pure, monomeric sample.

### Static light scattering measurement

Static light scattering (SLS) kinetic experiments were performed using a custom-built instrument equipped with a 475-nm laser source and 2-mm path-length quartz cuvettes. Depolymerization kinetics of hetero-O1 and hetero-O3 oligomers were monitored at 60 °C using oligomer concentrations of 1μM (expressed as monomer equivalents). This concentration was chosen to prevent repolymerization, as previously reported (20). Templating experiments were conducted by incubating purified uniformly ^13^C-labeled wt O1 oligomers with ^12^C-labeled I206A monomers, and the kinetics were monitored at 42°C by recording the time-dependent scattered light intensity at 475 nm. In these experiments, ^13^C-wt O1 oligomers were used at a concentration of 3μM (monomer equivalents) and I206A monomers at 30μM. Scattered light intensity was collected at a 90° angle and used as a proxy for changes in the weight-average molecular mass of species present in solution. Under Rayleigh scattering conditions, the scattered intensity I(t) is proportional to the sum of squared aggregation numbers weighted by species concentration, according to: *I(t) = I_0_.R.M Σ^n^_i=1_ C_i_i^2^*. Where I_0_ is the incident light intensity, R is the Rayleigh constant specific to the instrument and wavelength, M is the molecular weight of monomeric PrP, C_i_ is the concentration of species i, and *i* is its aggregation number.

### High resolution AFM imaging

High-resolution atomic force microscopy (AFM) experiments were performed using a NanoWizard 3 instrument (JPK Instruments) operated in quantitative imaging (QI) mode under liquid conditions. Measurements were carried out in 20 mM sodium citrate buffer (pH 3.5) using PeakForce HIRES FB cantilevers (Bruker) with a nominal spring constant of 0.12 N·m⁻¹ and a nominal tip radius of 10Å. For sample preparation, 50 μL of oligomeric assemblies were deposited onto freshly cleaved mica and incubated for 5 min. Protein concentrations were adjusted to 50 nM for purified O1 oligomers and to 150 nM (expressed as monomer equivalents) for the condensed O1 assemblies. After incubation, the mica surface was rinsed three times with imaging buffer to remove unbound material. AFM images were acquired in liquid by fixing the peak force set point at 90 pN.

### AFM-IR

AFM-IR measurements were performed using a NanoIR Icon instrument (Bruker Nano Surfaces, Santa Barbara, CA), in which pulsed infrared radiation is delivered from above the sample and focused near the AFM cantilever apex. Local infrared absorption induces photothermal expansion of the sample, detected through excitation of cantilever resonances. The instrument was coupled to a multichip quantum cascade laser (MIRcat, Daylight Solutions), providing tunable mid-infrared excitation over the 1800–900 cm⁻¹ range with 1 cm⁻¹ spectral resolution and pulse repetition rates up to 2 MHz. Measurements were carried out in tapping mode following established procedures (26). PPP-NCHAu-MB cantilevers (Nanosensors) were used, with the second eigenmode (∼1.6 MHz) sustaining the tapping oscillation and the first resonance (∼260 kHz) used for infrared signal detection. Samples were deposited onto freshly cleaved mica substrates using the same protocol as for high-resolution AFM imaging. After adsorption, samples were gently rinsed with deuterium oxide to shift the amide II band to the amide II’ region and reduce spectral contributions from residual water, then dried under a gentle nitrogen flow prior to analysis. Hyperspectral AFM-IR maps were acquired at 1590 cm⁻¹ and 1690 cm⁻¹ to selectively probe uniformly ¹³C-labeled wild-type PrP and ¹²C-labeled mutant PrP, respectively (see also SI-3). Normalized absorption signals were combined to generate composite overlay maps using Mountain Maps software (version 10.1, Digital Surf). Local AFM-IR spectra were acquired at defined positions on individual oligomeric assemblies, with four successive spectra averaged per location to improve signal-to-noise ratio. Spectral preprocessing included correction for QCL chip discontinuities followed by vector normalization, and analyses were performed using in-house Python scripts.

### ATR-FTIR

For secondary-structure analysis of hetero-O1 assemblies, hetero-oligomerization was performed by incubating 50 μM uniformly ¹³C-labeled wild-type PrP monomers with 100 μM uniformly ¹²C-labeled I206A monomers under oligomerization conditions. Following incubation, hetero-O1 oligomers were purified by size-exclusion chromatography as described above. The purified fraction was then concentrated to 30 μM (expressed as monomer equivalents) using MicroSpin concentrators prior to spectroscopic analysis. ATR-FTIR spectra were recorded using a Nicolet iS50 FTIR spectrometer (Thermo Fisher Scientific) equipped with a liquid-nitrogen-cooled mercury–cadmium–telluride (MCT) detector. Measurements were performed using a single-reflection germanium ATR crystal (Specac). Spectra were acquired at a resolution of 4 cm⁻¹ by co-adding 200 interferograms and analysed in the amide I region to resolve the isotopically shifted contributions of ¹³C- and ¹²C-labeled PrP.

### Kinetic simulations under arrested reaction conditions

ARC was modeled using a minimal kinetic framework describing a two-step oligomerization pathway comprising soluble monomers (M), a transient oligomeric intermediate (T), and a stable oligomeric species (O). The model incorporates concentration-dependent nucleation and growth steps, such that stable oligomer formation is kinetically suppressed at low monomer concentration while transient intermediates can still be populated within a finite observation time. The system of ordinary differential equations was implemented in MATLAB and numerically integrated (ode45) over a fixed time window for increasing initial monomer concentrations, with intermediate and oligomer species initially set to zero. Final species abundances were used to generate the concentration-dependent profiles shown in Fig. 5A, illustrating the ARC regime experimentally accessed by temperature-arrest protocols.

## Acknowledgements

This work has been financially supported by: French National institute for Agricultural Research (INRAe), French Medical Research Foundation (Equipe FRM DEQ20150331689) and the support of Hochschule Fresenius university (Germany). S.Prigent post-doc was founded by INRA-package for excellence and the European Research Council (ERC Starting Grant SKIPPERAD, number 306321).

## Author contributions

Conceptualization, H.R., DM, and SP.; Methodology, H.R., DM., A.I., SP., A.D.; J.M.; V.B.; G.v.R; S.L; J.T; H.K; A.D.B; AD; J.M; and P.S; Software, H.R.; Formal Analysis, HR; D.M; S.P.; Data Curation, HR and D.M.; Writing Original Draft, H.R.; D.M. and S.P.; Writing – Review & Editing, H.R.; D.M. and S.P; Visualization, H.R.; D.M.; and S.P.; Supervision, H.R. and D.M.

---

**SI-1.**
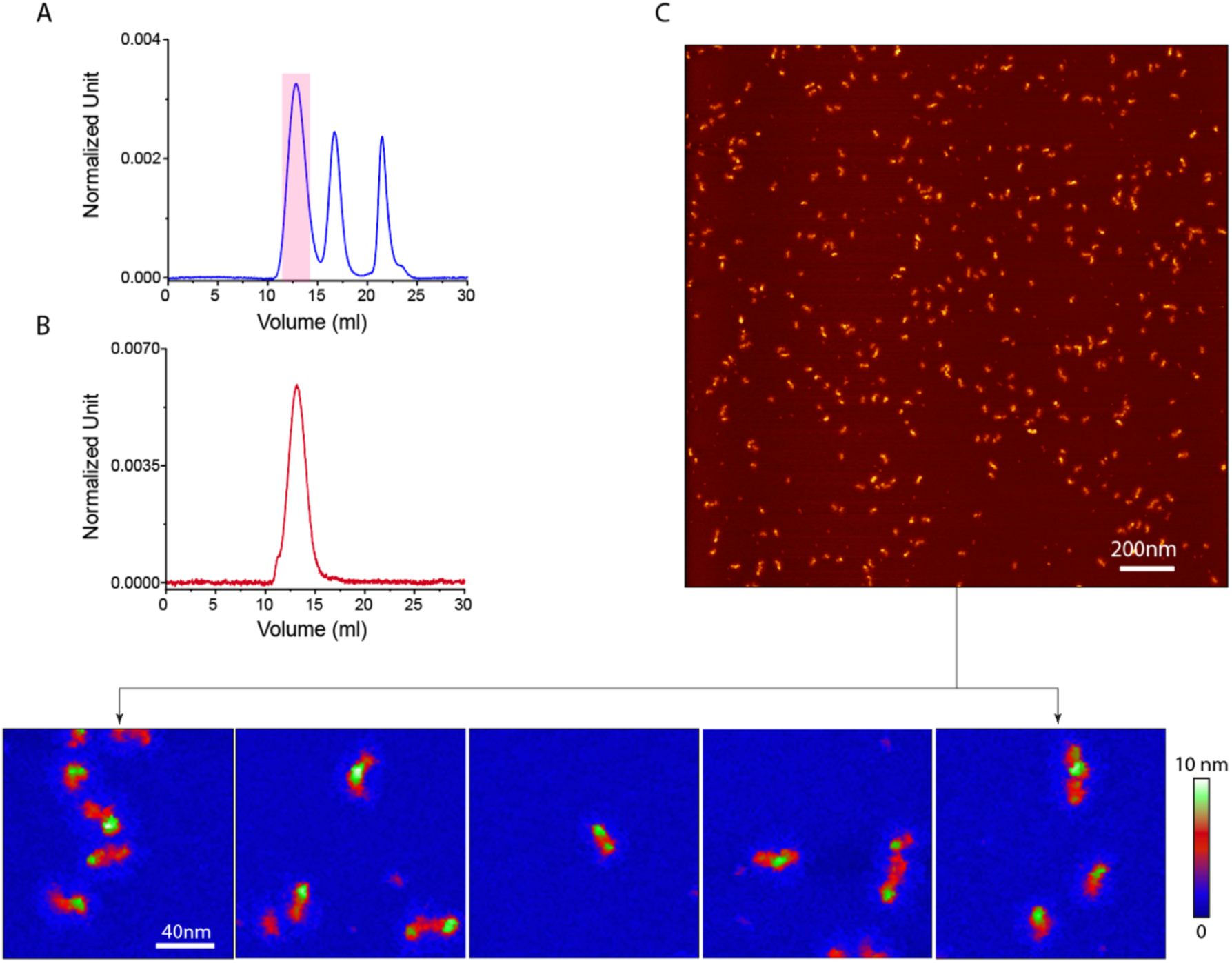

## Typical purification of O1 oligomers

As described previously, full-length ovine prion protein (PrP) variants subjected to thermal unfolding at pH values below 4.5 or above 6.0 can evolve into distinct soluble oligomeric species termed O1, O2, and O3 (refs). In the present study, O1 oligomers were generated by incubating full-length ovine PrP (A136, R154, Q171 variant) at 50 °C for 60 min under acidic conditions. Longer incubation times could be applied without detectable alteration of O1 assembly properties.

Following thermal unfolding, the reaction mixture was injected onto a TSK4000SW size-exclusion chromatography column, and the elution peak corresponding to O1 oligomers was collected as indicated in Fig. SI-1A. Purified O1 fractions were subsequently subjected to additional size-exclusion chromatography to assess oligomer stability and exclude depolymerization, and were further characterized using AFM based approaches.

Figure SI-1C shows representative AFM images of purified O1 oligomers, illustrating the heterogeneity of individual assemblies. The lower panel presents magnified views of selected regions from panel C, highlighting the structural features of individual O1 particles.

**SI-2.**
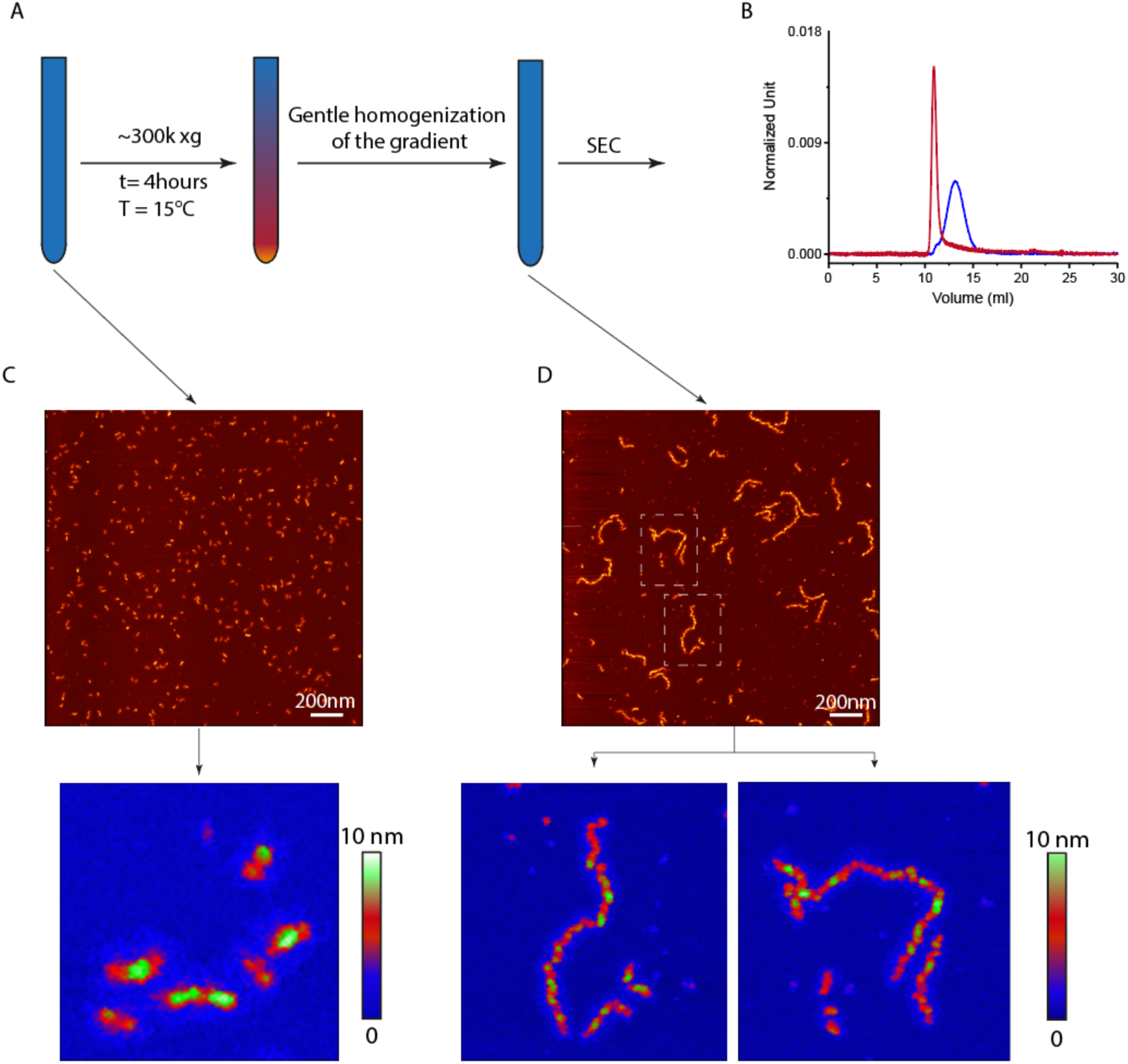

O1 oligomers have previously been reported to undergo condensation into larger supramolecular assemblies upon local concentration increase. In the present study, O1 condensation was induced by increasing the local concentration of SEC-purified O1 oligomers using an ultracentrifugation-based isopycnic concentration approach. As illustrated in Fig. SI-2A, purified O1 assemblies (blue SEC elution profile in Fig. SI-2B) were subjected to ultracentrifugation at 300,000 × g for 4 h at 15 °C, resulting in a marked increase in local oligomer concentration without altering buffer composition.

Following ultracentrifugation, the entire solution volume was gently homogenized to redistribute the material uniformly prior to further analysis. Importantly, this homogenization step did not lead to the reappearance of small O1 oligomers, indicating that the condensation process is irreversible under these conditions. Subsequent size-exclusion chromatography (Fig. SI-2B, red population) and high-resolution AFM analyses (Fig. SI-2C and D) revealed a pronounced shift toward larger assemblies after ultracentrifugation, consistent with stable formation of higher-order supramolecular structures derived from O1 oligomers. These observations demonstrate that O1 condensation is a concentration-driven but kinetically trapped process, leading to persistent supramolecular assemblies rather than a reversible equilibrium with smaller oligomeric species.

**SI-3.**
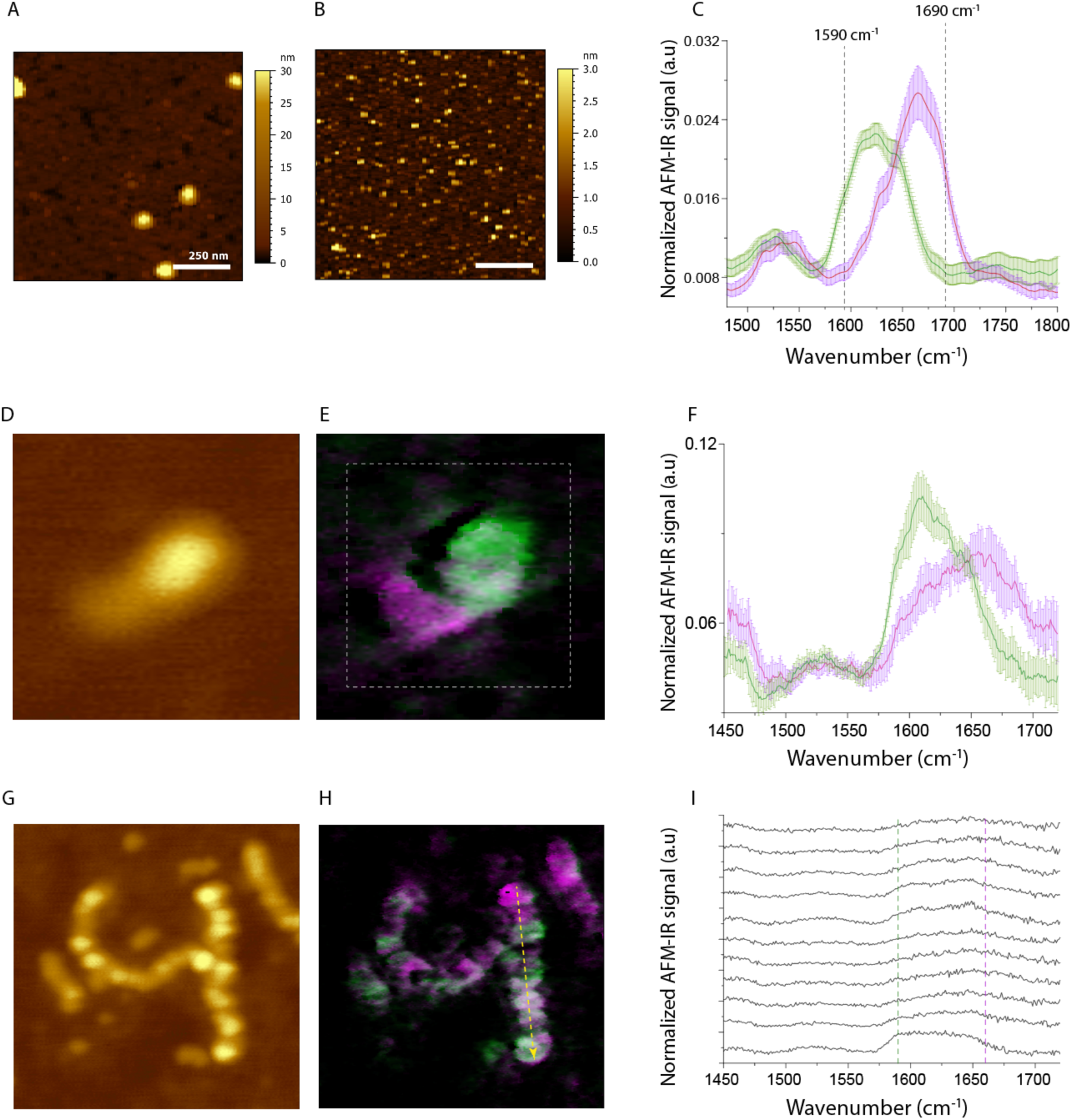

## Determination of reference wavenumbers for isotopically labeled PrP and validation of AFM-IR hyperspectral analysis

To establish reference wavenumbers enabling discrimination between ¹²C- and ¹³C-labeled PrP by AFM-IR, O1 oligomers generated from uniformly ¹²C-labeled wtPrP and uniformly ¹³C-labeled wtPrP were analyzed independently. Samples (50 μL at 1 μM) were deposited onto freshly cleaved mica substrates for 3 min, followed by three successive washes with 100 μL of D₂O to reduce spectral contributions from residual water and shift the amide II band to the amide II’ region. Representative low-resolution AFM topography images of ¹²C- and ¹³C-labeled O1 assemblies are shown in panels A and B, respectively. A total of 80 nano-IR spectra were recorded from ¹²C-labeled oligomers and 45 spectra from ¹³C-labeled oligomers. Mean averaged spectra are presented in panel C. The isotopic shift observed in the amide I region enabled identification of characteristic absorption maxima at approximately 1690 cm⁻¹ for ¹²C-labeled protein and 1590 cm⁻¹ for ¹³C-labeled protein. These wavenumbers were subsequently used as reference channels for selective detection of isotopic species in hyperspectral AFM-IR experiments throughout this study.

Typical hyperspectral acquisitions (panels D and E) were performed over square regions encompassing individual oligomeric assemblies using a step size of approximately 50 nm. Individual spectra were classified using k-means clustering (Quasar software), yielding two distinct spectral classes corresponding to substrate regions, ¹³C-labeled areas, and ¹²C-labeled areas. For the isotopically labelled classes, averaged spectra are shown as solid lines (green for ¹³C-labeled and purple for ¹²C-labeled), with shaded envelopes indicating the corresponding standard deviations.

Panels G and H illustrate representative AFM topography and corresponding isotopic localization maps of condensed hetero-O1 assemblies obtained after isopycnic concentration. The spatial distributions of ¹²C-labeled I206A (pink) and ¹³C-labeled wtPrP (green) are shown. Panel I presents a representative hyperspectral acquisition recorded along the trajectory indicated by the yellow arrow in panel H, illustrating spectral variations along the length of the condensed assembly.

## References

1. D. C. Bolton, M. P. McKinley, S. B. Prusiner, Identification of a protein that purifies with the scrapie prion. Science 218, 1309–1311 (1982).

2. M. L. Sprunger, M. E. Jackrel, Prion-Like Proteins in Phase Separation and Their Link to Disease. Biomolecules 11 (2021).

3. F. Chiti, C. M. Dobson, Protein Misfolding, Amyloid Formation, and Human Disease: A Summary of Progress Over the Last Decade. Annu Rev Biochem 86, 27–68 (2017).

4. O. V. Galzitskaya, S. O. Garbuzynskiy, M. Y. Lobanov, Prediction of amyloidogenic and disordered regions in protein chains. PLoS Comput Biol 2, e177 (2006).

5. L. D. Aubrey, S. E. Radford, How is the Amyloid Fold Built? Polymorphism and the Microscopic Mechanisms of Fibril Assembly. J Mol Biol 437, 169008 (2025).

6. H. Rezaei et al., Amyloidogenic unfolding intermediates differentiate sheep prion protein variants. J Mol Biol 322, 799–814 (2002).

7. H. Rezaei et al., Sequential generation of two structurally distinct ovine prion protein soluble oligomers displaying different biochemical reactivities. J Mol Biol 347, 665–679 (2005).

8. F. Eghiaian et al., Diversity in prion protein oligomerization pathways results from domain expansion as revealed by hydrogen/deuterium exchange and disulfide linkage. Proc Natl Acad Sci U S A 104, 7414–7419 (2007).

9. G. Spagnolli et al., Full atomistic model of prion structure and conversion. PLoS Pathog 15, e1007864 (2019).

10. A. W. Fitzpatrick et al., Atomic structure and hierarchical assembly of a cross-beta amyloid fibril. Proc Natl Acad Sci U S A 110, 5468–5473 (2013).

11. A. J. Marchut, C. K. Hall, Side-chain interactions determine amyloid formation by model polyglutamine peptides in molecular dynamics simulations. Biophys J 90, 4574–4584 (2006).

12. D. E. Koshland, Application of a Theory of Enzyme Specificity to Protein Synthesis. Proc Natl Acad Sci U S A 44, 98–104 (1958).

13. E. Chatani, N. Yamamoto, Recent progress on understanding the mechanisms of amyloid nucleation. Biophys Rev 10, 527–534 (2018).

14. V. N. Uversky, Mysterious oligomerization of the amyloidogenic proteins. FEBS J 277, 2940–2953 (2010).

15. D. M. Walsh et al., Amyloid beta-protein fibrillogenesis. Structure and biological activity of protofibrillar intermediates. J Biol Chem 274, 25945–25952 (1999).

16. J. R. Brender et al., Probing the sources of the apparent irreproducibility of amyloid formation: drastic changes in kinetics and a switch in mechanism due to micellelike oligomer formation at critical concentrations of IAPP. J Phys Chem B 119, 2886–2896 (2015).

17. M. Hoshi et al., Spherical aggregates of beta-amyloid (amylospheroid) show high neurotoxicity and activate tau protein kinase I/glycogen synthase kinase-3beta. Proc Natl Acad Sci U S A 100, 6370–6375 (2003).

18. X. Yu, J. Zheng, Polymorphic structures of Alzheimer’s beta-amyloid globulomers. PLoS One 6, e20575 (2011).

19. M. Adrover et al., Prion fibrillization is mediated by a native structural element that comprises helices H2 and H3. J Biol Chem 285, 21004–21012 (2010).

20. N. Chakroun et al., The oligomerization properties of prion protein are restricted to the H2H3 domain. FASEB J 24, 3222–3231 (2010).

21. Z. Xu, S. Prigent, J. P. Deslys, H. Rezaei, Dual conformation of H2H3 domain of prion protein in mammalian cells. J Biol Chem 286, 40060–40068 (2011).

22. C. Huin et al., Conformation-dependent membrane permeabilization by neurotoxic PrP oligomers: The role of the H2H3 oligomerization domain. Arch Biochem Biophys 692, 108517 (2020).

23. S. Simoneau et al., In vitro and in vivo neurotoxicity of prion protein oligomers. PLoS Pathog 3, e125 (2007).

24. F. Collu, E. Spiga, N. Chakroun, H. Rezaei, F. Fraternali, Probing the early stages of prion protein (PrP) aggregation with atomistic molecular dynamics simulations. Chem Commun (Camb) 54, 8007–8010 (2018).

25. H. Rezaei et al., High yield purification and physico-chemical properties of full-length recombinant allelic variants of sheep prion protein linked to scrapie susceptibility. Eur J Biochem 267, 2833–2839 (2000).

26. J. Mathurin et al., How to unravel the chemical structure and component localization of individual drug-loaded polymeric nanoparticles by using tapping AFM-IR†. Analyst DOI: 10.1039/c8an01239c (2019).

27. A. Armiento et al., The mechanism of monomer transfer between two structurally distinct PrP oligomers. PLoS One 12, e0180538 (2017).

